# Measuring human cerebral blood flow and brain function with fiber-based speckle contrast optical spectroscopy system

**DOI:** 10.1101/2023.04.08.535096

**Authors:** Byungchan Kim, Sharvari Zilpelwar, Edbert J. Sie, Francesco Marsili, Bernhard Zimmermann, David A. Boas, Xiaojun Cheng

## Abstract

Cerebral blood flow (CBF) is an important indicator of brain health and function. Diffuse correlation spectroscopy (DCS) is an optical technique that enables non-invasive and continuous bedside monitoring of human CBF. However, traditional DCS consisting of a few channels has relatively low signal-to-noise ratio (SNR), preventing measurements at long source detector separations (SDS) with increased sensitivity to cerebral rather than extracerebral blood flow. Here we developed a fiber-based speckle contrast optical spectroscopy (SCOS) system and the corresponding data analysis pipeline to measure CBF variations. We show that SCOS outperforms traditional DCS by more than an order of magnitude in SNR with comparable financial cost. We also demonstrated human brain function measurements during a cognitive task at an SDS of 33 *mm*. This technology will establish the foundation for devices that use spatial speckle statistics to non-invasively monitor human CBF, leading to a new functional neuroimaging approach for cognitive neuroscience.

## Introduction

Cerebral blood flow (CBF) is an important indicator of brain health as it regulates oxygen delivery to the brain and removes metabolic waste such as carbon dioxide. Alterations in CBF correlate with serious clinical conditions such as ischemic stroke^1, 2^, traumatic brain injury^3^, and Alzheimer’s disease^4, 5^. CBF also provides information about brain function^6–9^ as neural activation induces hemodynamic changes via neurovascular coupling^10^. Thus, monitoring CBF is important for cognitive neuroscience studies as well as clinical applications. Diffuse correlation spectroscopy (DCS) is an optical technique that measures human CBF from coherent light re-emitted from the tissue^11–15^. The blood flow index (BFi), a metric linearly correlated with underlying blood flow, is calculated from the decorrelation time of the autocorrelation function of the speckle intensity time course. It offers a convenient way to non-invasively and continuously monitor CBF at the bedside that cannot be accomplished with other techniques such as positron emission tomography and arterial spin labeling magnetic resonance imaging. However, traditional DCS systems suffer from a relatively low SNR, and the single-photon avalanche diode (SPAD) detectors used in these systems are generally expensive making it non-ideal for high density geometries covering large brain regions. Recently, several groups have attempted to improve DCS SNR by either imaging multiple speckles onto a SPAD array or improving the photon flux detection per speckle. For example, a recently published work on multi-speckle DCS with 1,024 parallel detection channels^9, 16^ showed promising improvements in SNR, and demonstrated human forehead CBF variations at a short source detector separation (SDS) of *ρ* = 15 mm. But at ρ= 15 mm, the sensitivity to the brain is low and not feasible for measuring brain function^17^. In another example, implementing interferometry has shown to improve DCS SNR by achieving shot noise performance^18, 19^, but at an expense of increased complexity of the system. Finally, using a longer wavelength of 1064 nm as the input light source has also been shown to raise DCS SNR by increasing photon flux, thanks to the higher maximum permissible exposure (MPE) and lower energy per photon than that of shorter wavelengths, but this method requires even more costly superconducting nanowire single-photon detectors^20^.

Another category of optical techniques to measure CBF is laser speckle contrast imaging (LSCI)^21–24^. Instead of analysing the temporal statistics i.e. auto-correlation function of the speckles as in DCS, LSCI exploits the spatial statistics by calculating the spatial contrast of the speckle intensity patterns measured within a certain camera exposure time. The speckle intensity patterns are obtained using relatively low-cost complementary metal–oxide–semiconductor (CMOS) cameras that can capture millions of speckles with millions of pixels to improve SNR as opposed to SPADs used in traditional DCS that utilize a few speckles. However, traditional LSCI has primarily been used to obtain two-dimensional images of superficial CBF with wide-field illumination, mostly for small animals such as mice with cranial windows or thinned skulls. Recently, a technique derived from LSCI named speckle contrast optical spectroscopy (SCOS) and its tomographic expansion speckle contrast optical tomography (SCOT) have been demonstrated for free-space imaging with larger SDSs, enabling non-invasive measurements of blood flow in deeper regions in phantoms, human arms and small animal brains^25–28^. However, generalizing free space techniques to human brain function measurements over a large area is challenging due to the presence of hair, susceptibility to motion due to limited range of focus, and the limited field of view of the camera. In addition, various noise sources will induce bias in the measured spatial contrast in SCOS, which presents challenges for quantifying blood flow changes at the low photon flux regime typically encountered for human CBF measurements^29^.

Here, we developed a fiber-based SCOS system capable of measuring human CBF and brain function non-invasively. We created and experimentally validated a data analysis pipeline to remove the bias in the contrast induced by noise. We verified experimentally that the current design of our fiber-based SCOS system using a scientific CMOS (sCMOS) camera outperforms traditional DCS systems by an order of magnitude higher SNR at a comparable cost. We performed human brain function measurements for the first time using SCOS during a mental subtraction task at ρ = 33 mm, providing sufficient sensitivity to CBF variations. This work opens a new avenue for future development of high-channel count and high-density optical devices (for tomography measurements) that can continuously monitor human CBF and brain function with unprecedented signal quality at large SDSs. The cost can be further reduced in the future by using lower cost CMOS cameras compared to the sCMOS camera used in this study. Continuous monitoring of CBF with high signal quality offers new opportunities for cognitive neurosciences and clinical applications.

## Results

Schematic of the fiber-based SCOS system, example speckle images, and the illustration of the data processing pipeline are shown in Fig. 1. Coherent light was launched through the multi-mode fiber onto the forehead and the diffusely remitted light was collected by a detector fiber bundle and subsequently imaged by a sCMOS camera (Fig. 1a). To populate the camera pixels, we imaged two rectangular fiber bundles at the same SDS onto the same camera. We show two example images of the fiber bundles focused and slightly defocused, as shown in Fig. 1b. For SCOS measurements, we kept the fiber bundles slightly defocused to improve the spatial uniformity of the image from the individual fibers in the bundle. The data analysis pipeline in Fig. 1c removed the bias in the raw speckle contrast, 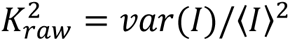, that arises from shot and instrumental noise sources and measurement non-uniformities, to obtain the fundamental spatial contrast squared 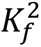 (see Methods), where *I* is the measured speckle intensity in the unit of camera counts (ADU). Here 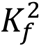 is directly related to the intensity auto-correlation function 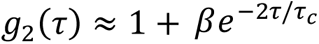 obtained from DCS measurements^30^, where τ is the delay time, τ_*c*_ is the decorrelation time, and β the coherence parameter. For long camera exposure times, *T*_*exp*_ ≫ τ_*c*_, which is valid in most human brain measurements 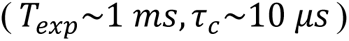, the blood flow index *BFi* is proportional 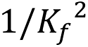^14, 22^.

**Fig. 1.**
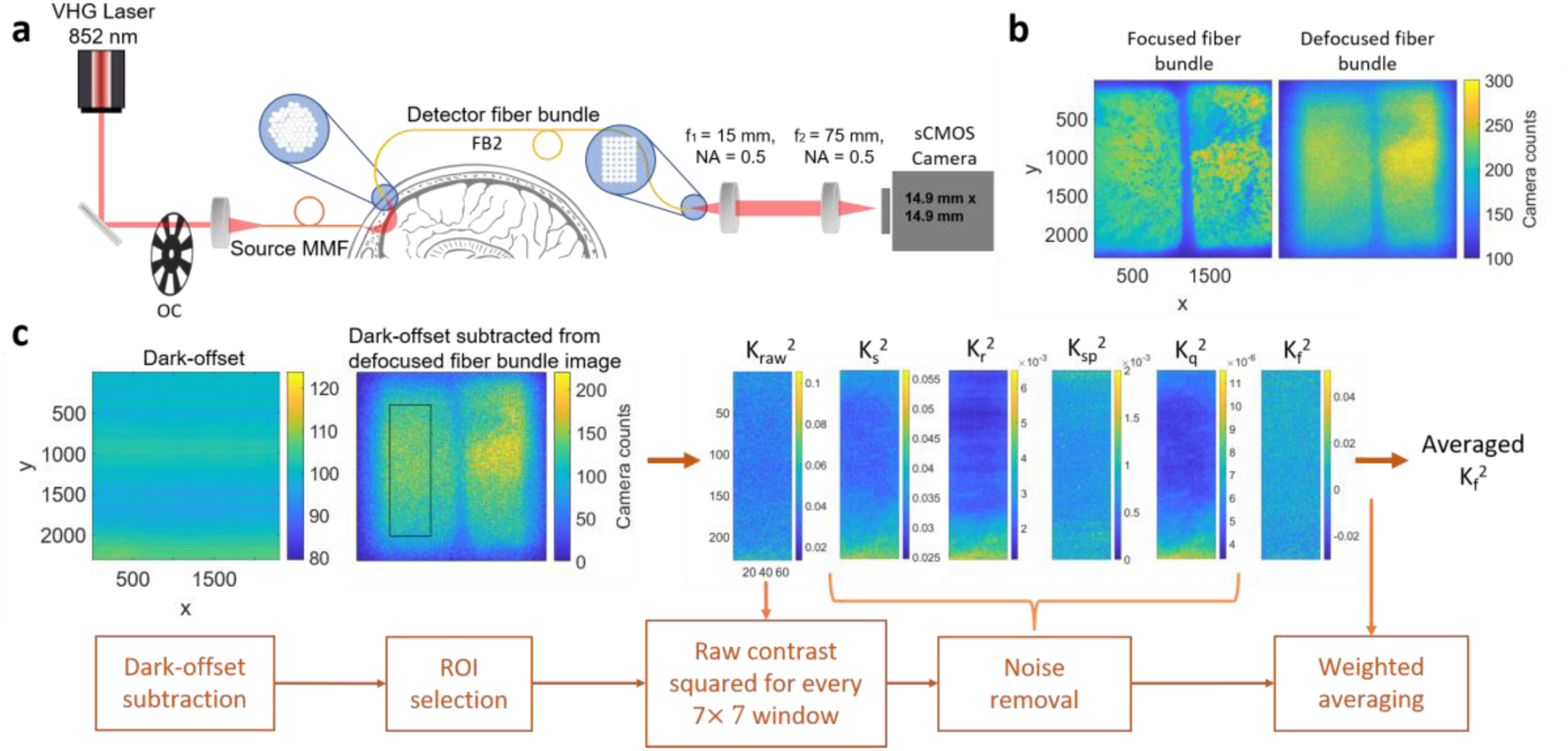
(a) The schematic of the fiber-based SCOS set-up. (b) Example speckle images when fiber bundles are focused and slightly defocused. (c) Illustration of the data analysis pipeline to remove the noise contributions from the measured 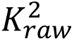 to obtain 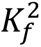 which is inversely proportional to *BFi.* VHG is volume holographic laser, OC is optical chopper, MMF is multi-mode fiber, FB is fiber bundle, NA is numerical aperture, CMOS is complementary metal-oxide-semiconductor, ROI is region of interest.

To validate the data processing stream, we tested SCOS measurements using a light-emitting diode (LED) light source (λ = 850 nm) as illustrated in Fig. 2. Since LED light is incoherent, no speckles will be generated and 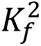 is expected to be zero. But 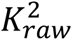 will be non-zero because of noise and non-uniformity in illumination. The measurement schematic and the results from measurements on a dynamic phantom sample are shown in Figs. 2a, c. We found that 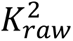 was approximately 0.02, reducing with increasing mean intensity because of reduced shot noise contribution to the bias, but that after correction 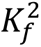 was greatly reduced towards the expected value of 0 (i.e. ∼10^−4^ − 10^−5^). The system schematic and results for human forehead baseline measurements at ρ = 33 mm are shown in Figs. 2b, d. We observed a cardiac pulse pattern in 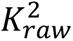 because of cardiac pulse induced changes in the measured intensity modulating the shot-noise contribution which modulates the measured contrast. After correction, the residual 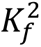 was on the order of 10^−5^, which is two orders of magnitude smaller than the blood flow induced 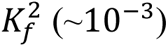 measured using coherent laser light in SCOS measurements. This illustrates the validity of the data processing stream to correct for the bias induced by noise in SCOS measurements.

**Fig. 2.**
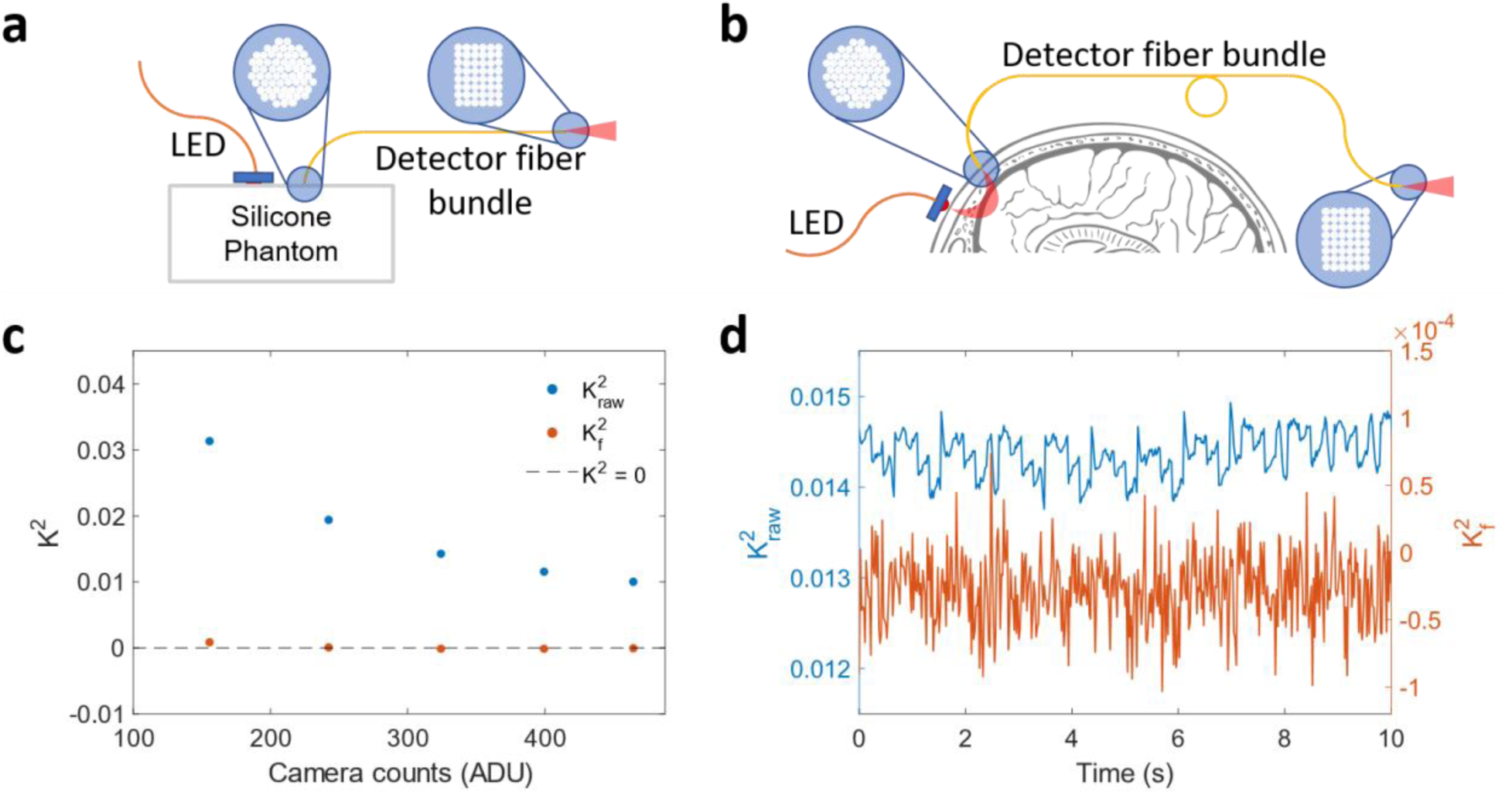
Validation of the data analysis pipeline for fiber-based SCOS. (a) Schematic experimental setup for LED measurements of the static phantom sample. The DC powered 850 nm LED is placed 15 mm away from detector fiber bundle. (b) Schematic experimental setup for human forehead LED measurements. (c) 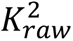 and 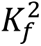 as a function of mean intensity 〈*I*〉 for the phantom sample LED measurements. (d) An example time course of 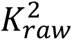 and 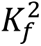 for human forehead LED measurements.

To improve the photon flux within a particular camera frame, we implemented a pulsing strategy with a rotating optical chopper (Fig. 1a). The duty cycle of the optical chopper was set at 10% so we could use 10x higher photon flux per camera exposure time while keeping the average incident power on the tissue within the maximum permissible exposure (MPE) of 33 mW, in compliance with ANSI safety standard limits at λ = 852 nm. This pulsing strategy should give an SNR gain by the square root of the photon flux increase, i.e. square root of 10x. We demonstrated the SNR gain by measuring cardiac signals with and without the pulsing strategy at ρ = 33 mm (Fig. 3a). The lower photon flux in CW mode resulted in higher instrumental noise, i.e. the measurement is in the read noise regime, leading to an inaccurate noise correction, likely because of instabilities in the camera read noise, and thus we obtained negative 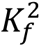 values. On the other hand, the higher photon flux in the pulsed mode overcomes the instrumental noise sources, permitting an accurate calibration to correctly estimate 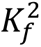 (see Supplementary Figs. 2-3 and discussion in the Supplementary for details). We observed an SNR improvement of ∼3x as expected using the pulsed mode (Fig. 3a). We also show examples of the measured cardiac signals at various SDSs ranging from ρ = 30 mm to 45 mm (Fig. 3b,c). At larger SDS, the correction for the bias became less accurate due to the instability of the read noise calibration for the camera we used. Thus, 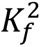 did not accurately report the absolute blood flow when read noise became appreciable at ρ = 40 mm and longer. Nevertheless, we still observed clear cardiac signal at ρ = 40 mm, indicating the possibility of using SCOS to measure relative blood flow changes at 40 mm or greater SDS provided a camera with a more stable read noise can be employed, or more speckles are integrated per camera pixel to maintain shot-noise performance.

**Fig. 3.**
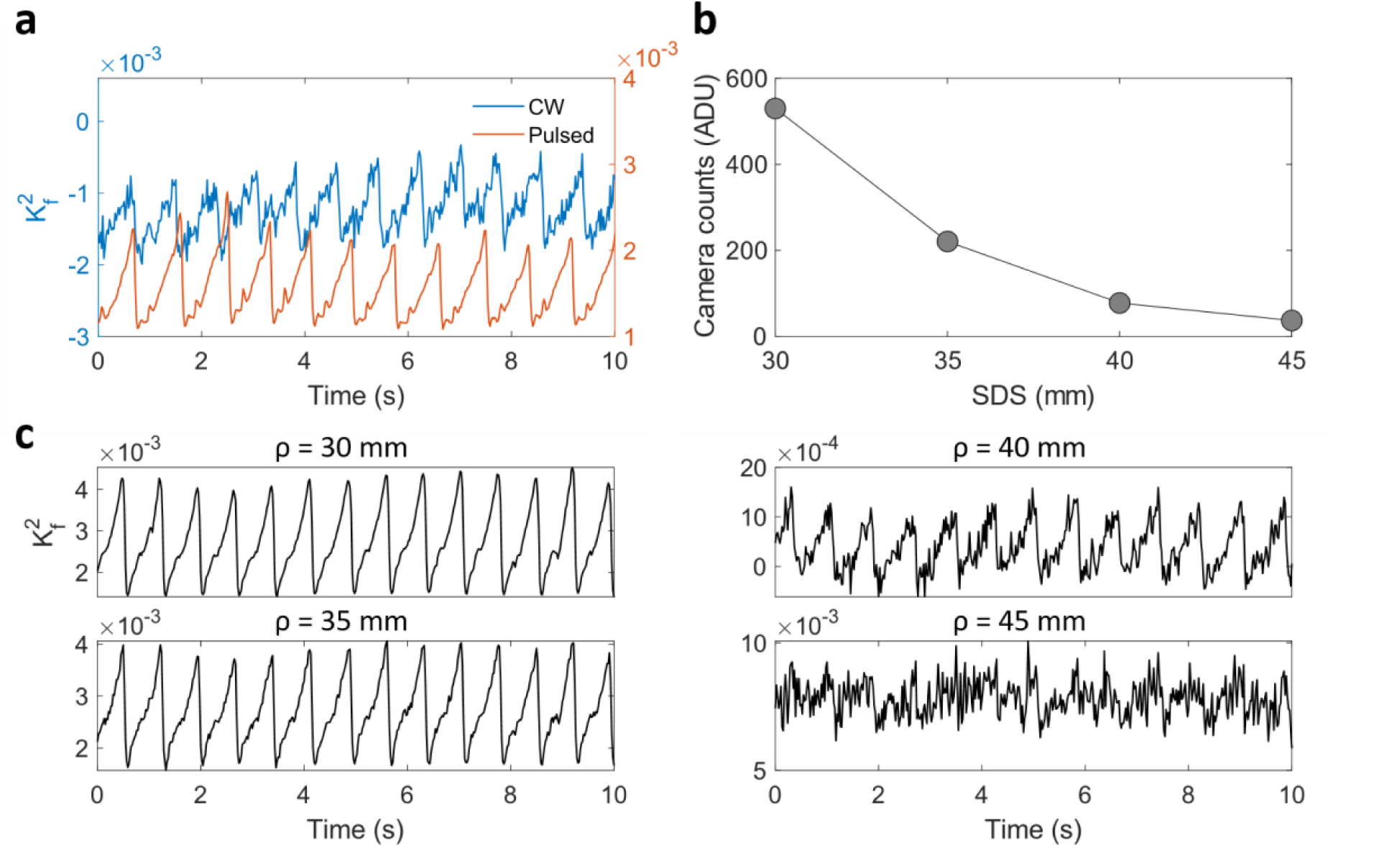
(a) Comparison of CW and pulsed fiber-based SCOS from baseline measurement on the human forehead at an ρ = 33 mm. (b) Mean camera counts and (c) time series of 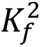 from baseline fiber-based SCOS measurements on the human forehead at SDS ranging from ρ = 30 to 45 mm.

Next, we compare the performance of SCOS with a state-of-the-art single-channel DCS system (Excelitas SPAD + Picoharp time tagger) using the same CW light source. We first compared the measurements using a liquid dynamic phantom (Intralipid 1.6% v/v in deionized water at room temperature) at ρ = 20 mm (Fig. 4a). Both the SCOS camera’s frame period and DCS g_2_(τ) integration time were set at 21.7 ms. 71.4% of the camera pixels were used for SCOS analysis, focusing on areas of high illumination from the fiber bundles. SNR was calculated as 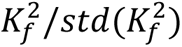 and 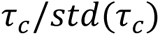 for SCOS and DCS measurements respectively. The SNR for DCS and SCOS was 5 and 114.8 respectively, representing a 23 fold increase afforded by SCOS. If the camera pixels were fully illuminated, we expect an 18% increase resulting in a 27 fold SNR improvement of SCOS versus DCS.

**Fig. 4.**
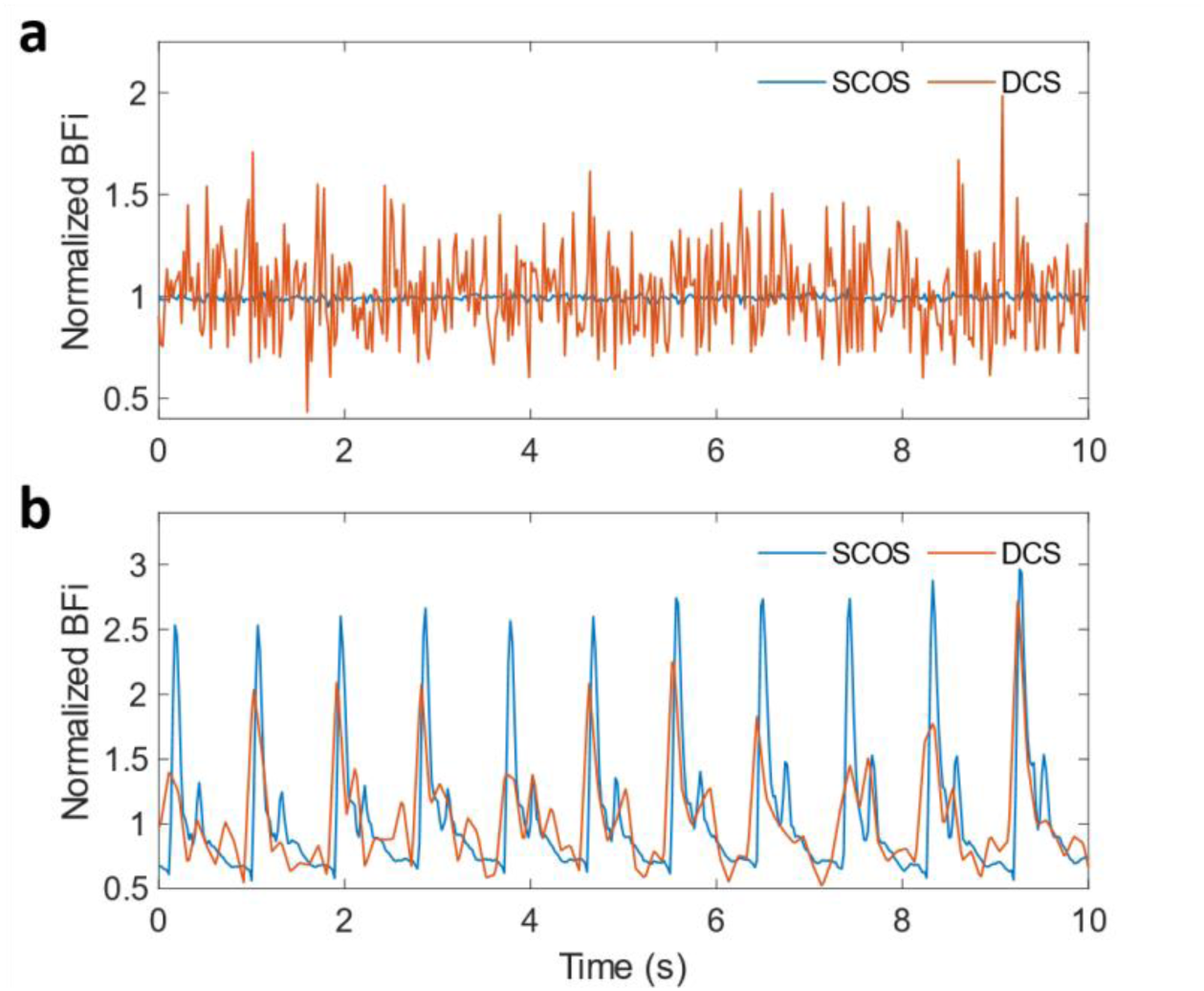
Comparison between fiber-based SCOS and DCS. (a) Normalized BFi of the dynamic phantom with 20mW CW at ρ = 20 mm. The frame period of fiber-based SCOS and the integration period of DCS are both set at 21.7 ms. (b) Human forehead normalized BFi with 35 mW CW at ρ = 20 mm. SCOS at 46 Hz (21.7 ms frame period) and DCS at 100 ms integration time.

We demonstrated this SCOS improvement in SNR over DCS qualitatively through measurements of cardiac signals on the human forehead using SCOS and DCS at a short SDS of ρ = 20 mm (Fig. 4b). Since the SNR for DCS is lower, we used significantly longer integration time of 100 ms for DCS to compare the signal quality with that of SCOS. Here *BFi* is calculated as 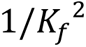 for SCOS and 1/τ_*c*_ for DCS. Despite the longer integration time for DCS, SCOS still has higher signal quality, which is apparent from the distinct features of the systolic peak and dicrotic notch in every pulse (Fig. 4b). This is consistent with the greater than order of magnitude difference in SNR observed with dynamic phantom samples.

In Fig. 5, we show the fiber-based SCOS measurement of blood flow and change in optical density (Δ*OD*) during a cuff-occlusion measurement on the human forearm (Fig. 5a). Δ*OD* is defined as Δ*OD* = *log*_10_(*I*_0_/*I*(*t*)), where *I*_0_ is the intensity at the baseline. When the applied external pressure is greater than systolic pressure, both the arteries and the veins were occluded. As expected, the *BFi* decreased during occlusion and there was a reduction in the ∼1 Hz cardiac fluctuations^20, 32^. The optical density increased since during the first few seconds of occlusion, the applied pressure only occludes the veins and arterial inflowing blood increases the blood flow, and thus total hemoglobin concentration. When the applied external pressure was released, both *BFi* and Δ*OD* overshot due to the classic hyperaemic overshoot to supply the oxygen starved tissue with oxygen, followed by a gradual recovery to baseline levels.

**Fig. 5.**
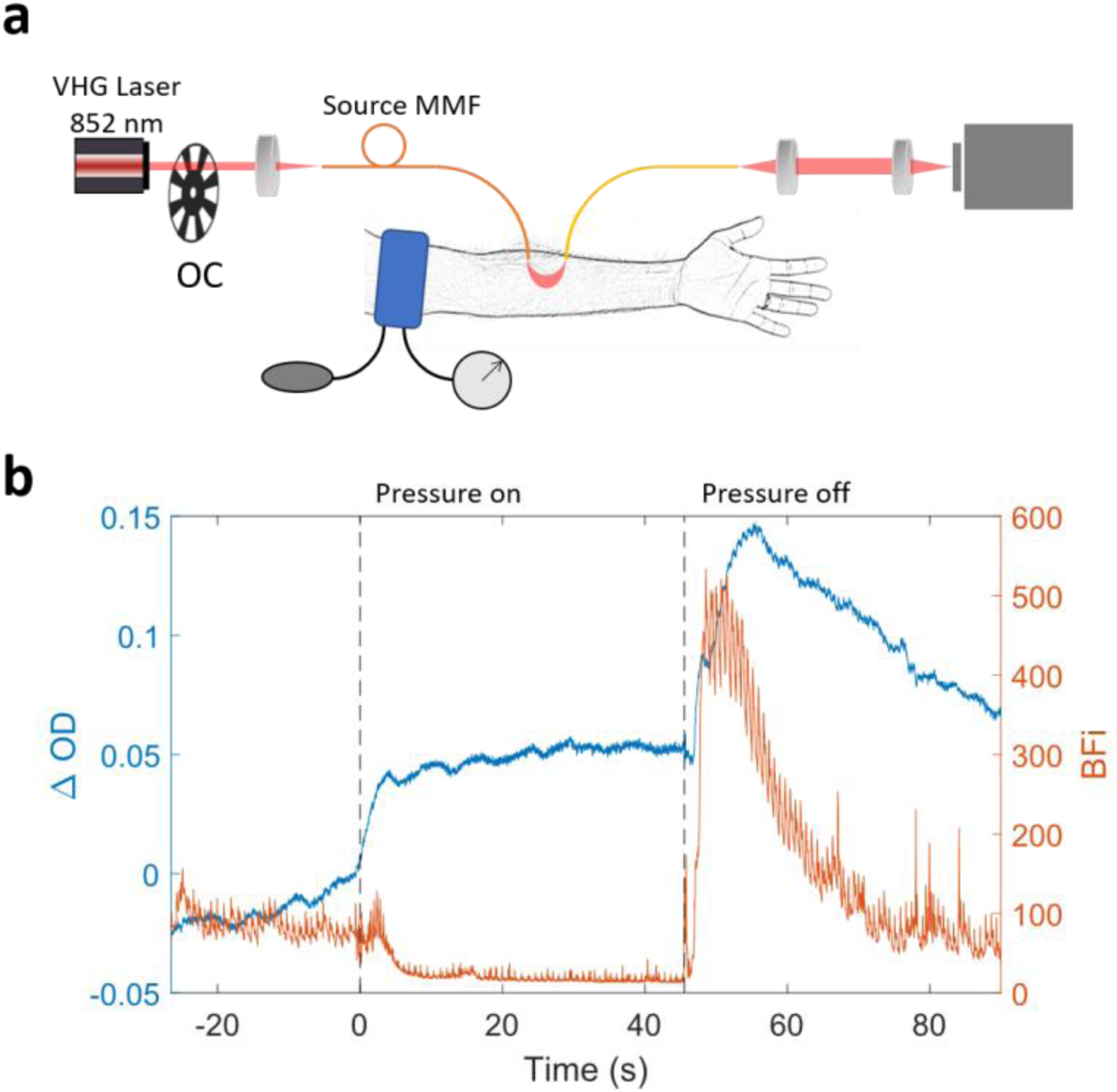
Cuff occlusion measurements using fiber-based SCOS. (a) Schematic representation of the cuff-occlusion measurements. (b) The time courses of BFi and Δ*OD* for cuff-occlusion measurements.

Finally, we show human brain function measurements during a mental subtraction task at ρ = 33 mm for three human subjects. As described in the Methods section, we have first used our existing high-density fNIRS system to locate the activation region on the forehead, and placed the fiber-based SCOS optodes at the location with largest response. Of the five subjects measured for SCOS, two did not show change in signal intensity (equivalent to single wavelength fNIRS) with mental subtraction. Since the same mental subtraction task (with different numbers) was used in the fNIRS measurements, it is possible that these subjects had already developed different strategies for the math problems thus not showing activation in SCOS measurements. We found that for all subjects with sufficient intensity changes during mental subtraction, blood flow increased by 7-16% during activation (Fig. 6c) and returned to baseline values post-activation, consistent with literature values^33, 34^. This work demonstrates the promising capabilities of SCOS for human brain function measurements.

**Fig. 6.**
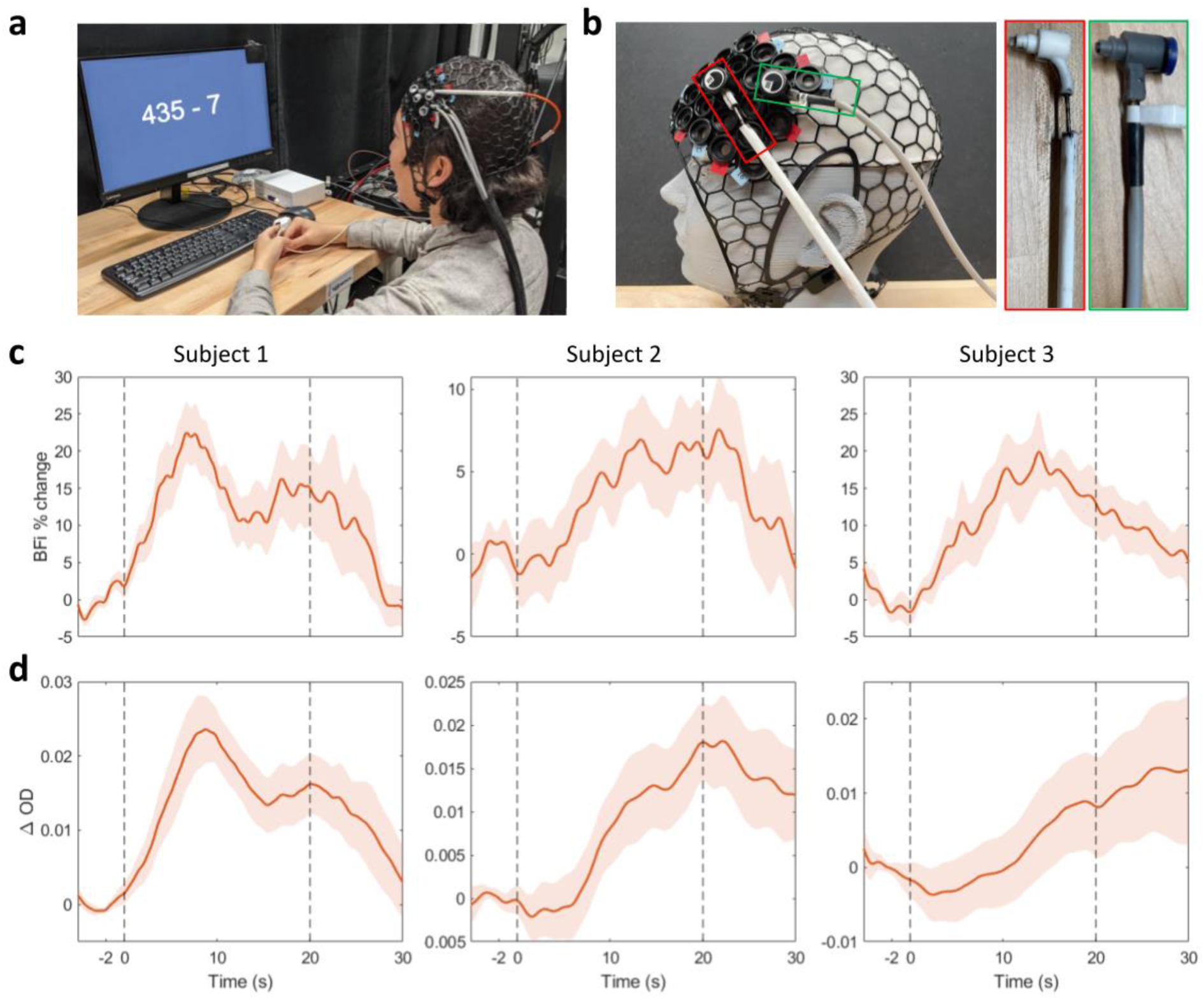
Mental subtraction measurements using fiber-based SCOS. (a) Example of the mental subtraction task conducted by a subject. (b) Placement of the NinjaCap and fibers on a subject head represented by the mannequin head. Source fiber (in red) and detector fiber bundle (in green) on a ninjaCap with NIRx spring loaded grommets. (c) Trial averaged BFi % change from baseline (t =-2 to 0 s) resulting from the mental subtraction task (presented from t = 0 s to t = 20 s. The shaded area indicates the standard error of the trial averaged response. (d) Trial averaged change in OD from baseline.

## Discussion

We have demonstrated for the first time a measurement of human CBF and brain function using a fiber-based SCOS system. We have developed and validated the data analysis pipeline to correct for noise-induced bias terms in the measured 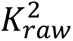 typically ignored in mouse brain LSCI measurements due to higher photon flux^20, 21^. With SCOS, shot-noise limited measurements can be achieved without employing interferometric detection, as has been done previously for DCS^19^ and SCOS^18^, by adjusting the measurement parameters including the speckle-to-pixel size ratio and the camera exposure time^29^. However, the integration of fiber-based SCOS (this work) with interferometry could be advantageous for measurements that desire a reduced camera integration time or have less photons per camera frame, or when using a camera with higher read noise. At the photon flux achieved with ρ = 33 mm on the human forehead, we believe there is little value in integrating interferometry to our fiber-based SCOS system.

In this manuscript, we have demonstrated the validity of our noise correction scheme for achieving shot-noise limited measurements. Using our recent SCOS noise model^29^, we found that we could achieve a shot noise limited measurement of speckle contrast when the camera count is above ∼100 ADU per pixel for our Hamamatsu camera. We used a pulsing strategy and low speckle-to-pixel ratio (s/p < 1) to achieve 200 ADU and greater camera counts for our human brain measurements at ρ = 33 mm (Fig. 2c). At lower camera count levels, our correction scheme was challenged mainly due to temporal variance in read noise for our camera (Supplementary Fig. 2). While keeping our measurements in the shot-noise limited regime, we improved noise correction by measuring dark images and estimating the read noise pattern before each measurement session. Additionally, by exploiting the use of a large number of pixels on the CMOS camera and utilizing the laser pulsing strategy, our fiber-based SCOS system can measure pulsatile *BFi* from the human forehead at up to ρ = 40 mm.

## Cost consideration

In the results section, we compared the performance of our fiber-based SCOS with a single channel DCS system. With cost taken into account, our fiber-based SCOS system (with an sCMOS camera) can achieve a 13.5x improvement in SNR of CBF estimation compared to a traditional DCS system (with a SPAD detector) with a similar cost (i.e. by increasing the DCS channel count to 4). The cost of a single channel SPAD detector >$3k while the cost of a sCMOS camera >$10k. For applications with less stringent SNR requirements, there are options for even lower cost CMOS cameras at an expense of higher read noise, lower bit depth, and potentially non-linear and non-uniform camera gain across pixels. For example, we carried out a preliminary measurement of the cardiac signal on the human forehead at ρ = 33 mm using a low-cost CMOS camera (Basler acA1920-160umPRO), which shows a promising high signal quality (Supplementary Fig. 4). While this is beyond the scope of the current manuscript, we believe it is important in the future to characterize different camera options that could be suitable and cost-effective for different applications of SCOS systems. Apart from the cost consideration, we found that the photon flux per speckle in our SCOS system is about 9 times smaller than that of the DCS. Some contributing factors include the energy loss in the lens system and the lower coupling efficiency of higher-order modes in multi-mode fibers. Future work could look into improving the optical design to narrow this gap to achieve even better performance from SCOS systems.

## Methods

### fiber-based SCOS system

The schematic of our fiber-based SCOS system is illustrated in Fig 1. The input laser light source (Thorlabs free-space VHG, 852 nm) is coupled into a multi-mode fiber (200 μm core diameter, 0.5 NA) which delivers the light to a sample such as the human forehead. We utilized custom-made rectangular shaped fiber bundles (∼1.64 x 3 mm, ∼3770 strands of 37 µm core diameter multimode fiber, 0.66 NA) as detectors and imaged two fiber bundles from different locations on the human scalp at the same SDS onto a single sCMOS (Hamamatsu, Orca Fusion BT) camera. The lenses form a 4f system as illustrated in Fig. 1a. The camera operates with the default settings of 16-bit resolution, fast scan operating mode with an expected read noise of 1.4 e^-^ across the sensor. We slightly defocus the imaged speckle to fill the gaps between individual fibers within a single fiber bundle as shown in Fig. 1b. The speckle to pixel size ratio (s/p) for the system has been calibrated to be 0.57 from 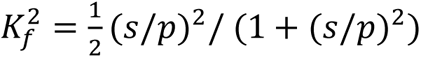 for unpolarized light obtained using a static phantom sample^29^ at high photon flux values to reduce errors introduced by noise.

## Data analysis pipeline for fiber-based SCOS

Raw data of intensity patterns for all the camera frames are recorded. The data processing stream is shown in Fig. 1c. We first subtract the raw intensity pattern by the dark offset which is obtained as the average of 500 dark images taken when the laser light is turned off. We then select the region of interest (ROI) for the speckle contrast calculation. Raw contrast squared is calculated as 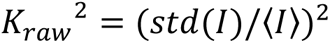 for each sliding window composed of 7 × 7 pixels within the ROI, where the intensity *I* is measured in ADU for all the pixels. We have used linear fitting to reduce the noise arising from estimation of 〈*I*〉 for each window. We first obtain the average intensity for all the pixels as a function of time denoted as *I*_*all*_ (*t*). We assume that the temporal shape of the average intensity for each window denoted as *I*_*w*_(*t*) is linearly related to *I*_*all*_(*t*) as *I*_*w*_(*t*) = *a* ∗ *I*_*all*_ (*t*) + *b*. The coefficients *a* and *b* are obtained from fitting and 〈*I*(*t*)〉 = *a* ∗ *I*_*all*_ (*t*) + *b*, which is a smoothed version of *I*_*w*_(*t*), is used as the denominator to calculate *K*_*raw*_ for each window. Examples of *I*_*all*_ (*t*), *I*_*w*_(*t*), and *a* ∗ *I*_*all*_ (*t*) + *b* are shown in the Supplementary Fig. 1. After obtaining the raw contrast squared *K*_*raw*_^2^, we remove the bias induced by shot noise 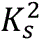, read noise 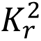, spatial non-uniformity in illumination 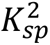, and quantization error 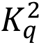 as

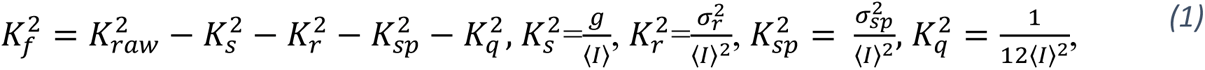

where *g* is the camera gain, σ_*r*_ is the read noise of the cameras, σ_*sp*_ is the variance obtained from the temporally averaged speckle image over 500 camera frames. 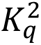 comes from the digital format of the camera output, in which a step size of one adds variance of 1/12^35^. To avoid erroneous impact of noise sources, the pixels with high read noise value i.e. σr > 10 and hot pixels of the camera are ignored. This correction process is also done for each 7 × 7 pixels. We then perform a weighted average of 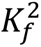 by *I*^2^ for all the windows within the ROI to obtain a single value of 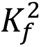 for a camera frame. We calculate 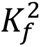 for all the camera frames to obtain the time course of 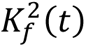. The average intensity *I*_*all*_ (*t*), simplified as *I*(*t*), is also obtained for all the camera frames. 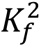 is related to the decorrelation time of τ_*c*_ and exposure time of *T*_*exp*_ via^21, 22^

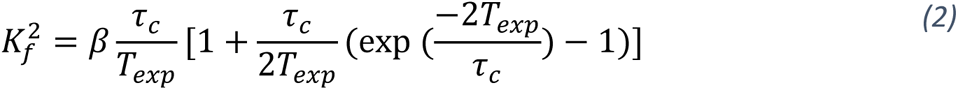

## Pulsing strategy to improve photon flux

We have utilized a pulsing strategy to improve the photon flux received within a camera frame. A rotating chopper wheel with 10% duty cycle is implemented to generate pulsed light to increase the photon flux within the camera exposure time while keeping the average incident power at 33 mW, within the ANSI safety limit at λ = 852 nm. The pulsed laser is synchronized with the camera frames during measurements by phase-locking the chopper wheel with the camera exposure sync signal, and as a safety precaution a 13 Hz trigger signal was utilized in place of the camera exposure sync signal when the camera was not exposing frames to ensure the light is pulsed while not performing measurements.

## DCS system and data analysis

We use a DCS system consisting of a single channel DCS (SPCM-AQ4C, Excelitas) with a Picoharp time tagger (PF300, PicoQuant). The source fiber and CW laser are shared with the fiber-based SCOS system. The DCS system is synchronized with the fiber-based SCOS for simultaneous data acquisition.

The intensity temporal autocorrelation function (g2(τ)) was generated from the raw intensity counts for time delay from 1 µs to 14.3 ms, with exponential increase in step between time delays. The bin size was set at 1 µs for the measurement time of 100 ms. The laser coherence (β) was estimated from fitting the g2(τ) curve averaged over the duration of the measurement to an exponential decay function given by^36, 37^:

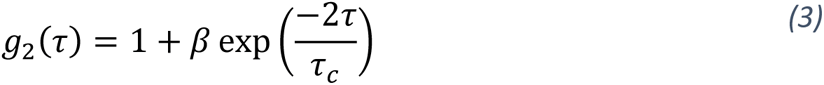

where τ is the time delay, and τ_*c*_is the decorrelation time. For each g2(τ) curve averaged over the measurement time the same exponential decay function and prior estimate of laser coherence is used to estimate the decorrelation time. The least-square curve fit was done using MATLAB’s lsqcurvefit function with implementation of the Levenberg-Marquardt algorithm.

## Dynamic phantom SNR measurement

The dynamic phantom was fabricated as 8% v/v ratio of 20% Intralipid (batch 10LI4282, Fresenius Kabi, Germany) in deionized water at room temperature. A 3D-printed mount made of flexible material (NinjaFlex TPU filament, NinjaTek) was used to mount the fiber-based SCOS fiber bundle and DCS single mode fiber at ρ = 20 mm from the source fiber.

## fNIRS and fiber-based SCOS brain activation data acquisition

To better localize the brain activation region, we utilize the high-density fNIRS system with a commercially available NIRSport2 (NIRx) with 14 sources and 32 detectors on the prefrontal cortex region with first and second nearest neighbour separation distances of 19 and 33 mm respectively. The sources and detectors are placed in a NinjaCap, a in-house 3D printed fNIRS cap made of flexible material (NinjaFlex, NinjaTek) with NIRx grommets for compatibility with variable tension spring holders. The cap is placed on the head with respect to EEG 10-10 midline central electrode site (Cz), which is estimated through measurement of nasion, inion, left/right pre-auricular points. The signal quality for each source-detector pair is tested through the Aurora fNIRS acquisition software (NIRx). fNIRS measurement during mental subtraction task is also done using the Aurora fNIRS acquisition software. The fNIRS data is analyzed with Homer3^38^ to obtain trial averaged oxyhemoglobin (HbO), deoxyhemoglobin (HbR), and total hemoglobin (HbT) changes for all source-detector pairs. The source-detector with largest HbO and HbT change is selected for subsequent fiber-based SCOS measurement.

For SCOS measurements, the same NinjaCap used in fNIRS measurements is placed on the subject’s head, followed by placement of custom source fiber and detector fiber bundle at the pre-selected source-detector pair. After baseline measurement for confirmation of photon flux, the subject undergoes mental subtraction task. The externally triggered camera sends images via Camera Link cable and card to the computer where the images are written to hard drive for processing.

## fNIRS data analysis

The fNIRS data is analysed using a processing stream in Homer3. The processing stream consists of (1) pruning channels with a SNR threshold of 10, (2) converting intensity values to change in OD, (3) applying a motion detection and correction algorithm^39^, (4) bandpass filtering the data from 0.01 to 0.5 Hz to remove signal drift and cardiac fluctuations, (5) converting change in OD to changes in HbO and HbR concentration using modified Beer-lambert law^40^, and (6) applying a general linear model (GLM) with signal from ρ =19 mm.

## Cuff-occlusion and mental subtraction

A sphygmomanometer with an analog manometer and a manually operated bulb was used to apply cuff occlusion. The inflatable cuff was placed around the left arm. The fiber-based SCOS source and detector fibers were placed on the left ventral forearm at ρ = 30 mm. The subject was optically measured for 120 s, consisting of 30 s of baseline, 45 s of cuff occlusion, and 45 s of return to baseline. For cuff occlusion, the bulb was squeezed repeatedly to increase the pressure to 180-200 mmHg within five seconds. The pressure was kept above 180 mmHg until pressure release happening over less than five seconds (Fig. 5).

During the mental subtraction task, the subjects are seated in front of a computer monitor providing mental subtraction task stimulus. After the baseline measurement, the subject is visually presented with a random selection of a three-digit number and a smaller number (6, 7, or 13) in a mathematical expression format (e.g. 270 – 7). The subject then serially subtracts the smaller number from the larger number (e.g. 270, 263, 256) until the problem disappears from the screen. Each problem is displayed for 20 s and the interval between problems is randomly selected from 10-20 s. Each measurement consists of five trials, with three measurements totalling 15 trials per subject for a total duration of 12 min.

## Participants

Nine subjects were recruited for this study, including seven subjects for measurement of mental subtraction induced CBF changes, one subject for the cuff occlusion measurement, and one subject for comparing cardiac signals between DCS and fiber-based SCOS, with and without pulsing strategy, and fiber-based SCOS over a range of SDSs. Of the seven subjects for mental subtraction task, five showed lateral frontal lobe activation and thus were subsequently imaged for fiber-based SCOS measurements. The experimental procedure and protocols were approved and carried out in accordance with the regulations of Institutional Review Board of the Boston University. Each subject provided a signed written informed consent form prior to the experiment.

## Data and code availability

The data and code that support the findings of this study are available from the corresponding author on reasonable request.

## Acknowledgements

This research was supported by funding provided by Meta Platforms Inc. The authors would like to thank Parya Farzam, Jessie Anderson, and De’Ja Rogers for help with using fNIRS, Meryem Yucel and Yuanyuan Gao for aid in fNIRS data processing, Joe O’Brien for aiding the development of source and detector probes for SCOS, and Mark Spatz for development of a software to acquire images from the SPAD and CMOS cameras.

## Author contribution

The presented idea is conceived from discussions among D.B., E.S., F. M., S.Z. and X.C.. S.Z. and X.C. developed the theory behind the noise correction and performed the computational modelling. B.K., B.Z., S.Z., and X.C. designed the optical setup and worked out the technical details. B.K., S.Z., and X.C. carried out the experiments, and contributed to the analysis and interpretation of results. D.B., E.S., F. M. and X.C. oversaw overall direction and planning. X.C. wrote the manuscript with support from all the co-authors. All authors read and agreed on the content of the paper.

## Supplemental Information

Supplementary Figures 1–4.

