## Supplementary Information for "Measuring human cerebral blood flow and brain function with fiber-based speckle contrast optical spectroscopy system"

### Linear estimation of window $\langle I(t)\rangle$ in the contrast measurement

As mentioned in the Methods section, we have calculated the contrast ${K_{raw}}^2 = (std(I)/\langle I\rangle)^2$ within each $7 \times 7$ window. To reduce the noise in $\langle I\rangle$, we have smoothed the average intensity within each window as described in the Methods. Examples of $I_{all}(t)$, $I_w(t)$, and $a * I_{all}(t) + b$ are shown in Fig. 1.

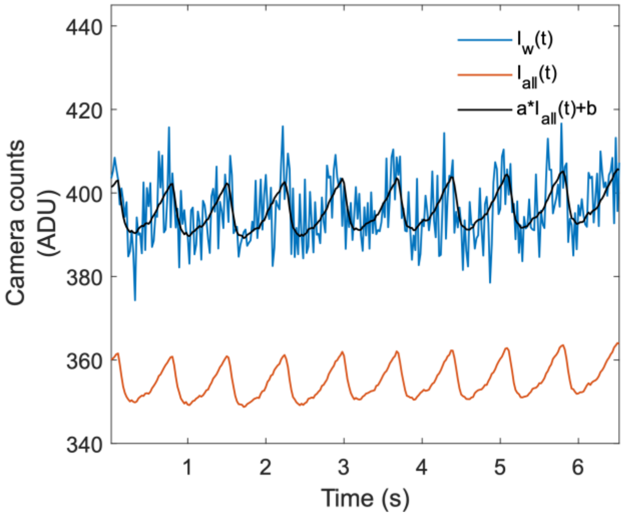


**Fig. 1**. Estimation of window mean intensity with linear fitting. Time series of mean camera counts for a 7 x 7 pixel window (blue) and for a 500 x 1500 pixel region of interest (ROI, red). The ROI time series is linearly fitted to the window time series for a less noisy estimate of window mean intensity time series (black).

### Dark images to obtain read noise and dark offset

We show the distribution of dark offset and read noise for all the pixels obtained from the standard deviation of the dark images on two different days in Fig. 2. We see that the read noise distribution slightly differ and this variation can induce variations of $K_r^2$ on the order of $10^{-3}$ (similar to $K_f^2$)

for ~100 ADU. To minimize the error, we always measure the dark images before an experiment and use the dark offset and read noise for the particular measurement session.

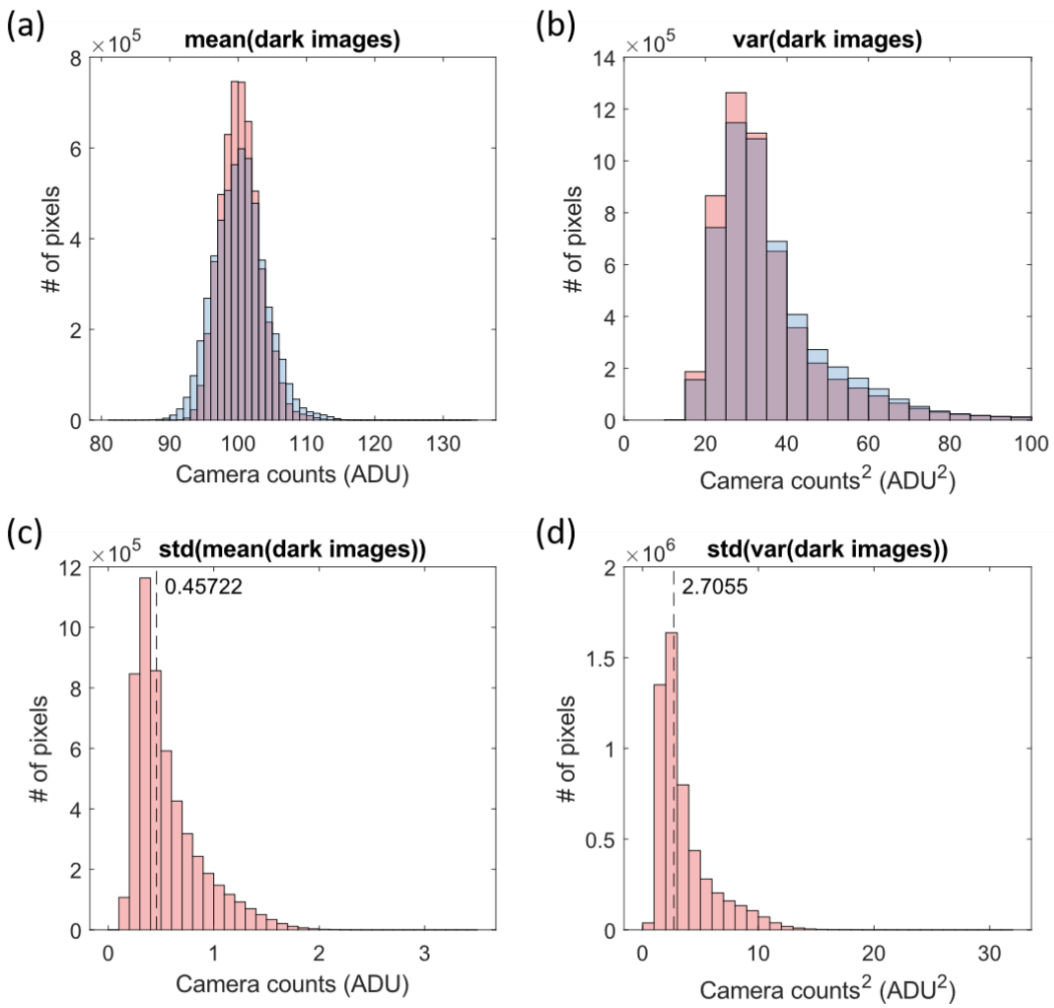


**Fig. 2**. Histogram of the (a) dark offset obtained from the average of 500 dark images and (b) read noise squared ($\sigma_r^2$) obtained from the variance of 500 dark images taken 10 minutes apart. Histogram of the (c) standard deviation of the pixel-by-pixel dark offset and (d) standard deviation of the pixel-by-pixel variance of dark images across multiple measurements on same day (n = 13).

#### The shot noise and read noise regime

We have numerically calculated the contribution of shot noise $K_s^2$ and read noise contribution $K_r^2$ to $K_{raw}^2$ as functions of camera counts (ADU) matching the experimental parameters described in the Methods section, using our previously established SCOS noise model in Fig.3. We see that for signals <100 ADU, the contribution of $K_s^2$ is smaller than $K_r^2$ (i.e. the measurement is in the read noise regime), making the noise correction scheme more susceptible to temporal instabilities in read noise shown in Fig. 2 and outputting inaccurate estimation of $K_f^2$. For human brain function measurements, the camera counts are ~200 ADU, where $K_s^2$ dominates (i.e. the shot noise regime) and the noise correction scheme is robust. Also, we see that for the shot noise contribution to become negligible, i.e., $K_{raw}^2 \sim K_f^2$, that the camera counts need to be $> 10^4$ ADU, which in general is not achievable for human brain measurements. This further confirms that the noise correction is necessary for human brain measurements using SCOS.

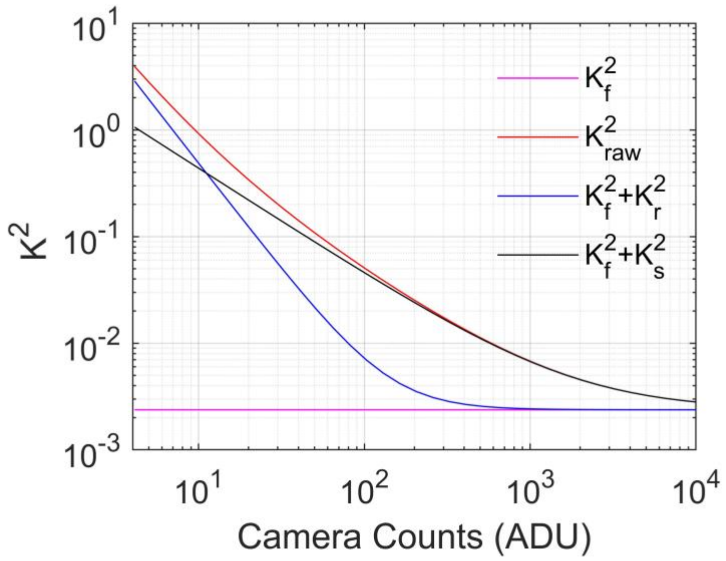


**Fig. 3**. $K_s^2$, $K_r^2$, $K_{raw}^2$, $K_f^2$ as functions of camera counts (ADU) with the parameters the same as in the experiments described in the Method section.

### Preliminary Basler cardiac measurements

We made a preliminary investigation into the use of a more cost-effective CMOS camera (Basler a2A1920-160umPRO) by measuring baseline cardiac signal with the same pulsed laser source set-up at $\rho$ = 33 mm in Fig.4. The camera was operated with 8-bit depth resolution, a frame rate of 46 Hz with estimated read noise of 1.6 e- and gain of 0.37 ADU/e-.

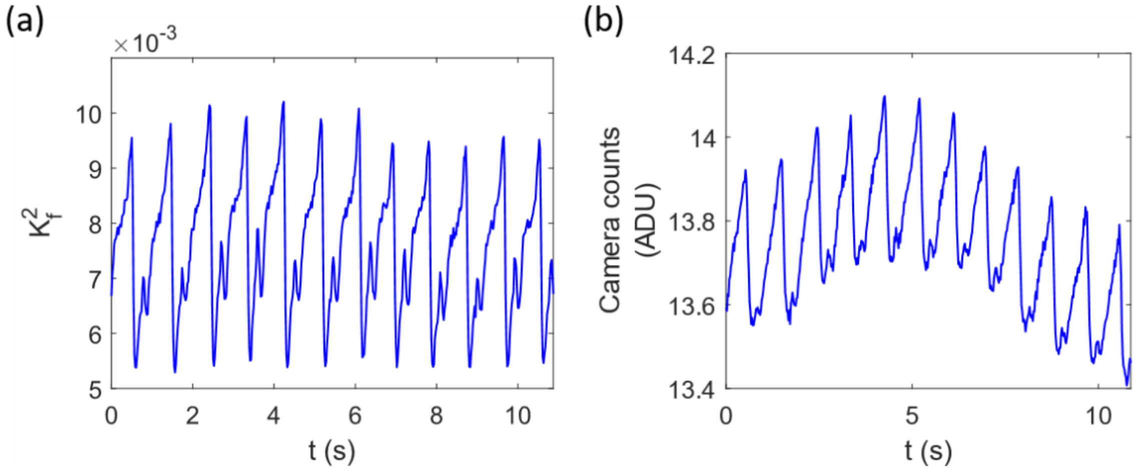


**Fig. 4**. (a) Fundamental contrast squared and (b) camera counts from the Basler camera baseline human forehead measurement at $\rho$ = 33 mm showing cardiac fluctuations.
